# Single-cell Spatial Explorer: Easy exploration of spatial and multimodal transcriptomics

**DOI:** 10.1101/2022.08.04.502890

**Authors:** Frédéric Pont, Juan Pablo Cerapio, Pauline Gravelle, Laetitia Ligat, Carine Valle, Emeline Sarot, Marion Perrier, Frédéric Lopez, Camille Laurent, Jean Jacques Fournié, Marie Tosolini

**Affiliations:** Université de Toulouse, Inserm, CNRS, Université Toulouse III-Paul Sabatier, Centre de Recherches en Cancérologie de Toulouse, Toulouse, France; ERL 5294 CNRS, Toulouse, France

**Keywords:** spatial transcriptomics, single-cell, multimodal analysis, visualization, open-source, freeware

## Abstract

The development of single cell technologies yields large datasets of informations as diverse and multimodal as transcriptomes, immunophenotypes, and spatial position from tissue sections in the so-called ‘spatial transcriptomics’. Currently however, user-friendly, powerful, and free algorithmic tools for straightforward analysis of spatial transcriptomic datasets are scarce. Here, we introduce Single-Cell Spatial Explorer, an open-source software for multimodal exploration of spatial transcriptomics, examplified with 6 human and murine tissues datasets.

## Introduction

By positioning in tissues the cells characterized by single cell RNA sequencing (scRNAseq), spatial transcriptomics (ST) unveils information beyond gene expression profiles and revolutionizes biology and medicine. Today, many packages using R or Python can map cells, score gene signatures, infer cell interactions, and detect spatial patterns [16]. However, computationally frugal visualization of multimodal ST images raises challenges not addressed so far. Most ST platforms [4, 8, 11, 12, 15] yield images of million cells and gigabytes of data which multimodal visualization require overlays with thousands cell scores for thousands gene signatures. So despite the current tools for pre-processing ST data and computing scores, producing such multimodal images remains challenging for (1) the joint availability of spatial coordinates and signature scores, (2) the immediate and tunable overlay of image and signature, (3) the user-friendly environment requested by biomedical users. We addressed comparable needs for non-ST scRNAseq datasets with Single-Cell Signature Explorer which, using a mere laptop, projects single cell’s signature scores across UMAP or t-SNE and proposes a slide mode visualisation of entire databases [6]. Other algorithms, whether command line-based or not also allow analysis of scRNAseq and CITE-seq datasets [1, 3, 13, 17]. Single-Cell Virtual Cytometer allows such analyses using laptops without command lines, and its inter-operability with Single-Cell Signature Explorer enables both phenotyping and cell sorting from numerical data [2, 7]. Since it was not possible to explore likewise ST datasets, here we introduce Single-Cell Spatial Explorer (scSpatial Explorer), a user-friendly and open-source software.

## Materials and Methods

### 0.1 Tissue samples

One human spleen sample was selected from the Centre de Resources Biologiques (CRB) collection at IUCT Oncopole CHU Toulouse (DC-2009-989) with a transfer agreement (AC-2008-820 Toulouse) obtained after approval from the appropriate ethics committees upon informed consent of the donor.

### 0.2 Staining and imaging and Spatial Transcriptomics

Snap-frozen human spleen samples were embedded in Tissue-Tek O.C.T. (Sakura). 10*µm*-thick slices were performed and placed on the Visium spatial slide (10x Genomics, Visium Spatial Protocols - Tissue Preparation Guide - Rev A), fixed with methanol, and stained with hematoxylin and aqueous eosin (HE). This HE-stained tissue was imaged with a ZEISS inverted microscope under 10x/0.3 objective, permeabilized, and reverse transcription was performed on the same slide. Additional examples of ST datasets for human cerebellum, spinal cord and murine brain and kidney were downloaded from the 10x Genomics website.

Second strand synthesis, denaturation, DNA amplification, libraries construction were performed following manufacturer’s instructions (10x Genomics, Visium Spatial Gene Expression Reagent Kits User Guide - Rev C). The librairies were profiled with the HS NGS kit for the Fragment Analyzer (Agilent Technologies) and quantified using the KAPA library quantification kit (Roche Diagnostics). The librairies were pooled and sequenced on the Illumina NextSeq550 instrument using a High Output 150 cycles kit and the cycles parameters : read 1 : 28 cycles, index 1 : 10 cycles, index 2 : 10 cycles, read 2 : 90 cycles.

### 0.3 Data preprocessing

Both of the HE-stained tissue image and the FASTQ data were then processed using SpaceRanger1.3.0 (10xGenomics) to link the image to the scRNAseq data. Prior to their analysis with scSpatial Explorer, the sized image of the sample and its corresponding transcriptome data had to be prepared as follows. First, using a scale factor and image size parameters (”scalefactors json.json” file), the tissue image and spatial array grid were superimposed to build a sized tissue image (Figure 2, top left). Using Seurat 4.1, transcriptomic data were normalized and exported as table.

**Figure 1.**
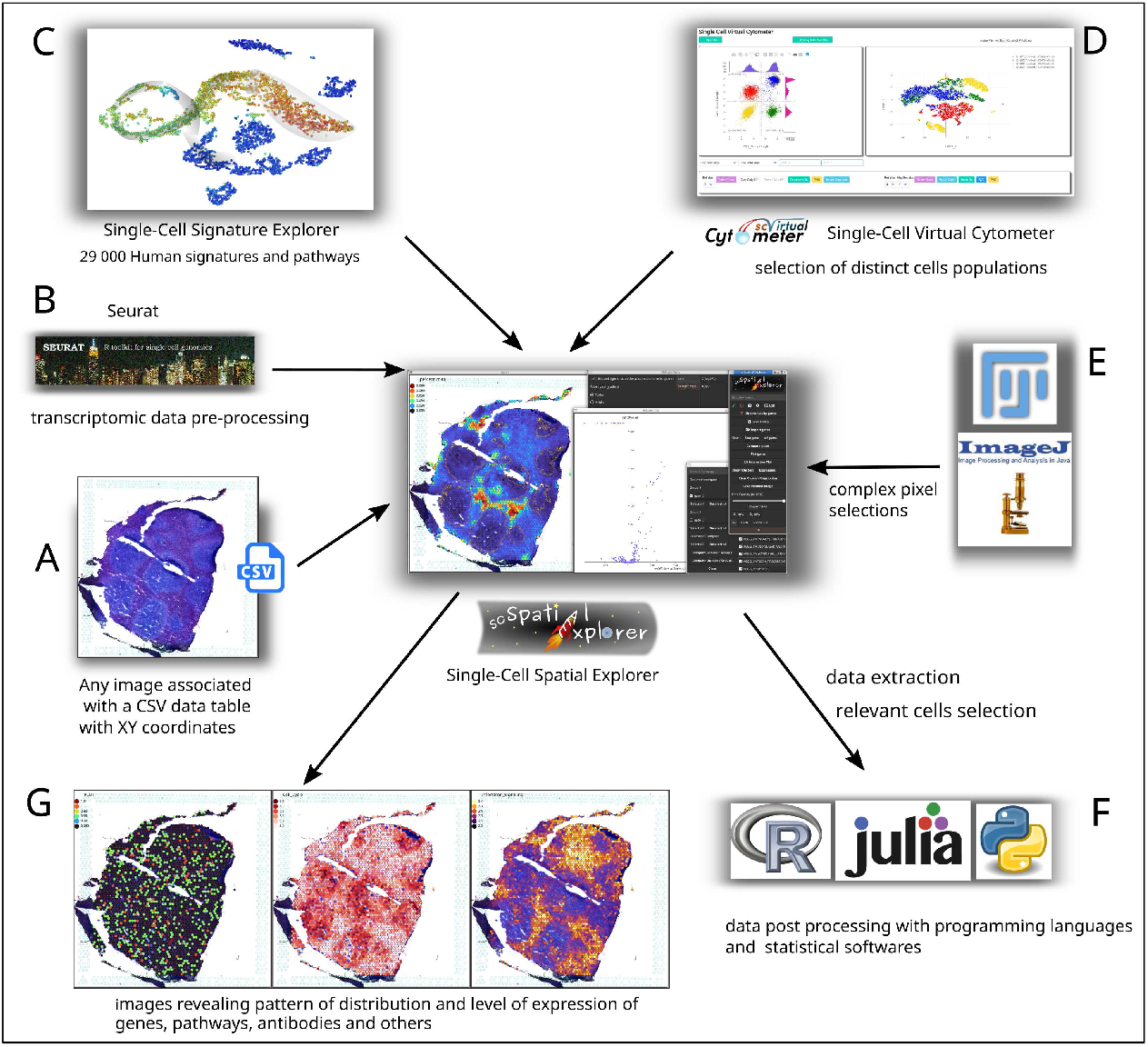
scSpatial Explorer interoperability with any PNG image associated to a table containing cells’ coordinates and other numeric data (A). ST pre-processed data, e.g. by Seurat (B) can be imported together with signature scores [6] (C), to be overlaid with the image. scSpatial Explorer is compatible with Single-cell Virtual Cytometer [7] for cells selections (D), it can import/export gates with ImageJ/Fiji format (E), and filter an unlimited number of tables from user-defined gates for further processing (F). scSpatial Explorer overlays the image with gene or antibodies expressions, or any numerical data. Color and opacity gradients tools improve scSpatial Explorer overlays (G) of image with any numerical data.

**Figure 2.**
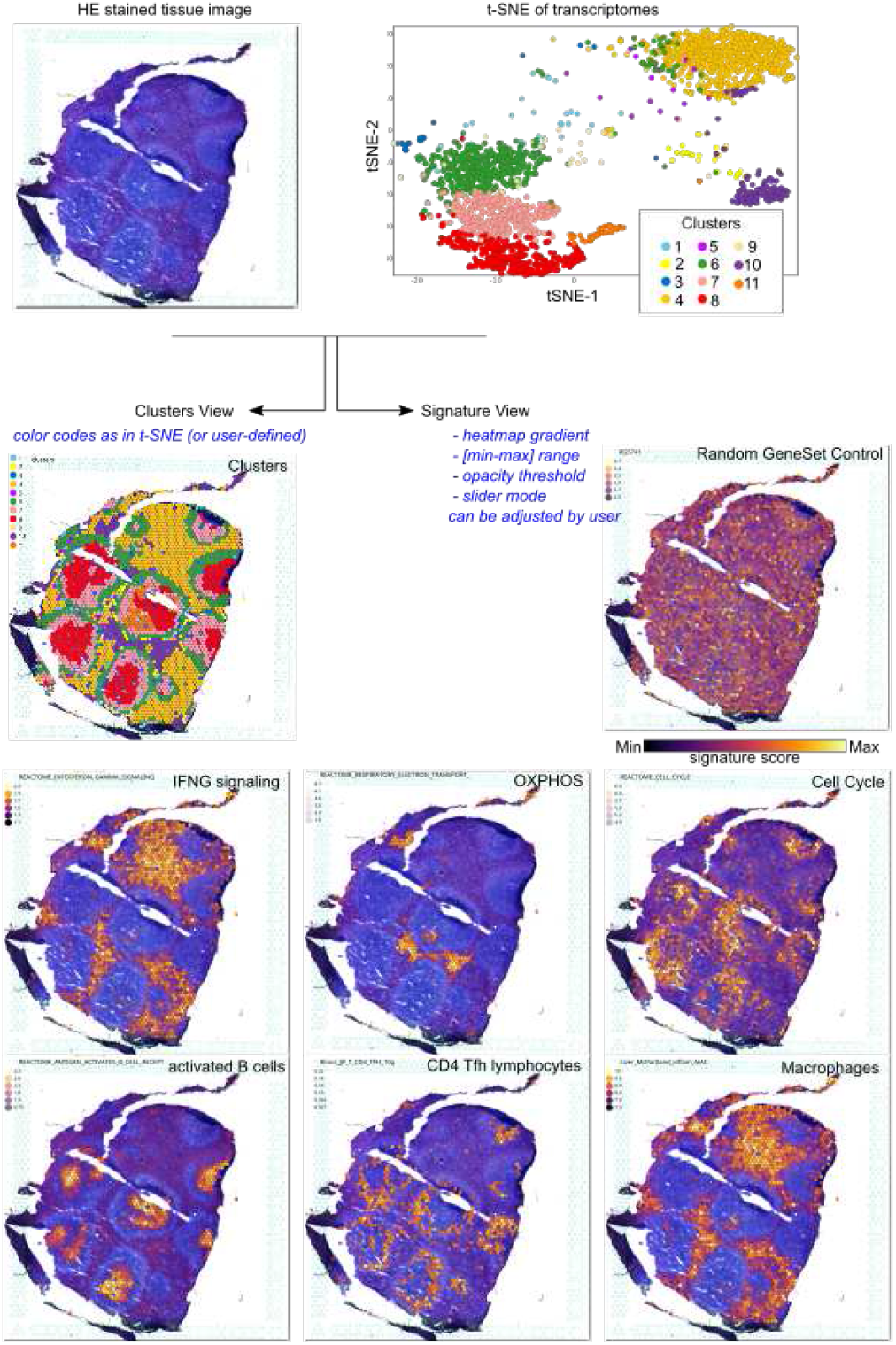
scSpatial Explorer pipeline. Cluster View (middle-left) overlays the tissue image (top-left) with the corresponding transcriptome clusters (top-right). The Signature View mode (middle-right) overlays the tissue image with the spot’s enrichment scores for a selected gene signature (‘Inferno’ gradient heatmap).

### 0.4 Code availability

scSpatial Explorer was developed in pure [Go](https://golang.org/) using the graphical library [Fyne](https://github.com/fyne-io/fyne). Precompiled static binaries are [available](https://github.com/FredPont/spatial/releases) for Linux, Mac and Windows. Files can be accessed at [GitHub scSpatial Explorer web page](https://github.com/FredPont/spatial).

## 1 Results and discussion

### 1.1 Single-Cell Spatial Explorer

The software is [easy to install](https://youtu.be/LBBkN_rJHuc) and can be run without any programming skills on any computer, and comes with a detailed [documentation](https://github.com/FredPont/spatial/blob/main/doc/Manual_Single_Cell_Spatial_Explorer.pdf) and [video tutorials](https://github.com/FredPont/spatial#demo-videos). It overlays on a microscopy image either the quantitative data (e.g. ST tables) shown as color gradient heatmaps or qualitative data (e.g. cluster or categories) delineated using discrete color palettes. It allows to gate any cell population delineated either by a region of interest (ROI) from the image or by a scatterplot. An unlimited number of data tables containing XY image coordinates can be [filtered by gates](https://youtu.be/rSefd0pmc7g), and such gates can be compared or plotted with any XY coordinates. Gate comparison displays an interactive volcano plot (Fold-Change *v*.*s*. P-value) which dots can be selected to get the corresponding expression data (see supplementary fig. 1).

scSpatial Explorer has been developed with inter-operability as priority, especially with Seurat [14], Single-Cell Signature Explorer [6] and Single-Cell Virtual Cytometer [7], leading to a very versatile and powerful solution (Fig. 1, Supplementary discussion ‘Features’). scSpatial Explorer is compatible with any PNG image and any data table (TAB separated files) containing XY image coordinates. Thus, the user can apply scSpatial Explorer at any level of his own analytic pipeline. For example, after raw data pre-processing, scSpatial Explorer can import gene or antibody expressions, as well as pre-computed signature scores (e.g. from MSIgDB [5, 6]). Cell populations gated with Single-Cell Virtual Cytometer [7] can also be imported for a flow cytometry-like analysis and unlimited sub-gatings. scSpatial Explorer can leverage ImageJ/Fiji image processing capabilities for image analysis, by macro-programming and gates importation [9, 10]. Sub-tables, cell names and gates coordinates can be [exported](https://youtu.be/rSefd0pmc7g) in CSV format for further processing.

Comparison with existing ST visualization softwares (Supplemental data) positioned scSpatial Explorer as a unique user-friendly tool.

### 1.2 ST analysis of a human spleen tissue

Below we exemplified ST analysis of a human spleen tissue produced by Visium technology (10X Genomics). The transcriptome data were visualized by t-SNE (Fig. 2, top-right) and the enrichment of all MSIgDB signatures were scored. Then, distinct exploration modes are available with scSpatial Explorer.

The ‘Cluster view’ mode overlays the sized tissue image and transcriptomic clusters delineated by a color palette. Here, the overlay of the spleen image with 11 transcriptomic clusters revealed a follicular pattern yet visible on the stained tissue, and the strictly follicular localisation of Clusters 6,7,8 (Fig. 2). The ‘Expression view’ mode overlays the tissue image with genes/antibodies expressions or enrichment scores. The user selects any signature from a large collection of curated or user-defined signatures. Negative control signatures (randomly selected genes) yield homogeneous images without pattern whereas curated signatures produced highly informative images (Fig. 2). Here, the follicular areas appeared strongly mitotic while their periphery expressed high IFN-signalling and mitochondrial respiration. Lineage-specific genesets identified antigen-activated B lymphocytes maturing in the follicles, peripheral rings of follicular helper CD4 T cells, and a macrophage-rich stroma (Fig. 2), in a spatial organisation reflecting the antibody-producing function of splenic germinal centers. In its ‘slider’ expression mode, scSpatial Explorer allows rapid screens of collections of signatures. Here, it pinpointed among others: germinal center, follicular plasmablasts, T lymphocytes, natural killers, dendritic cells, myeloid cells, Hallmarks such as ‘G2M checkpoint ‘, ‘Unfolded Protein Response’, and Transcription factor targets of CEBP2, HSD17B8, SKIL, and ZNF407.

scSpatial Explorer harnesses the image processing developments. To characterize the molecular hallmarks of spatially-defined cells, it can import gates with the same CSV polygon format as ImageJ/Fiji Region Of Interest (ROI) (see Supplementary information). Any cell-delineating pattern defined by image analysis can be imported likewise for analysis by scSpatial Explorer. Gates can also be directly drawn on the image using the lasso or polygon tools. Here, the transcriptomes from an intra-follicular gate and a peri-follicular gate were compared (Supplementary fig. 1). This comparison returns a Fold Change and P-value table, and an interactive volcano plot for the table columns selected by the user. Then, selection of any dot displays expression of the corresponding signature as a heatmap overlaid with the image. Here, the human splenic germinal centers up-regulated several signatures such as ‘CD22-mediated BCR regulation’ and ‘Mitosis’ reflecting their role in BCR-mediated selection of antigen-specific B lymphocytes.

Beyond identifying the transcriptome hallmarks of spatially-delineated cells/spots, scSpatial Explorer indeed allows to localize any image cells/spots selected by transcriptome criteria. For example, from a human prostate cancer ST [dataset](https://www.10xgenomics.com/resources/datasets/human-prostate-cancer-adjacent-normal-section-with-if-staining-ffpe-1-standard-1-3-0), cells gated in a scatterplot of “Hypoxia VS Androgen Response” signatures are instantly mapped within the IF image (Supplementary fig. 2). Such gates can also be delineated from scatterplots of single gene expressions, immunophenotypes, or any other numerical values from multimodal datasets.

To illustrate further how scSpatial Explorer performs with any kind of tissue colored by HE or immunofluorescence (IF), the following examples downloaded from [10x Genomics website](https://www.10xgenomics.com/) were analysed for diverse multigene signatures of cell lineages, metabolic and molecular pathways, as well as behavioural hallmarks from the Gene Ontology collection of the MSIgDB [5] database:

- human prostate cancer (IF) Supplementary Figure 2
- human Cerebellus (HE) Supplementary Figure 4
- human spinal cord (IF) Supplementary Figure 5
- mouse kidney (HE) Supplementary Figure 6
- mouse brain (HE) Supplementary Figure 7

## 2 Conclusion

Concluding, scSpatial Explorer is a very powerful, versatile, and interoperable tool for ST analysis. It rapidly overlays the tissue image with quantitative metrics, such as multigene signature scores. It harnesses a memory-efficient approach (Supplementary fig. 3) to the visualization needs of a large audience of biomedical scientists without advanced analytic and programming skills, packages, libraries, and wrappers. We forecast that when reaching the true single cell resolution level, the next generation of ST will benefit even more of scSpatial Explorer and its inter-operability, notably with emerging frameworks of spatial pattern meta-analyses.

## Acknowledgments

[Andrew Williams](https://andy.xyz), creator of the [Fyne project](https://github.com/fyne-io/fyne) and CEO at [Fyne Labs](https://fynelabs.com/), is acknowledged for his useful technical advice about the usage of the Fyne graphical toolkit.

This work was granted access to the HPC resources of CALMIP super-computing center under the allocation P19043. We are also grateful to the Génotoul bioinformatics platform (Bioinfo Genotoul, Toulouse Midi-Pyrenées) for providing computing resources.

## Supplementary Information

### Supplementary figures and table

**Supplementary Figure 1.**
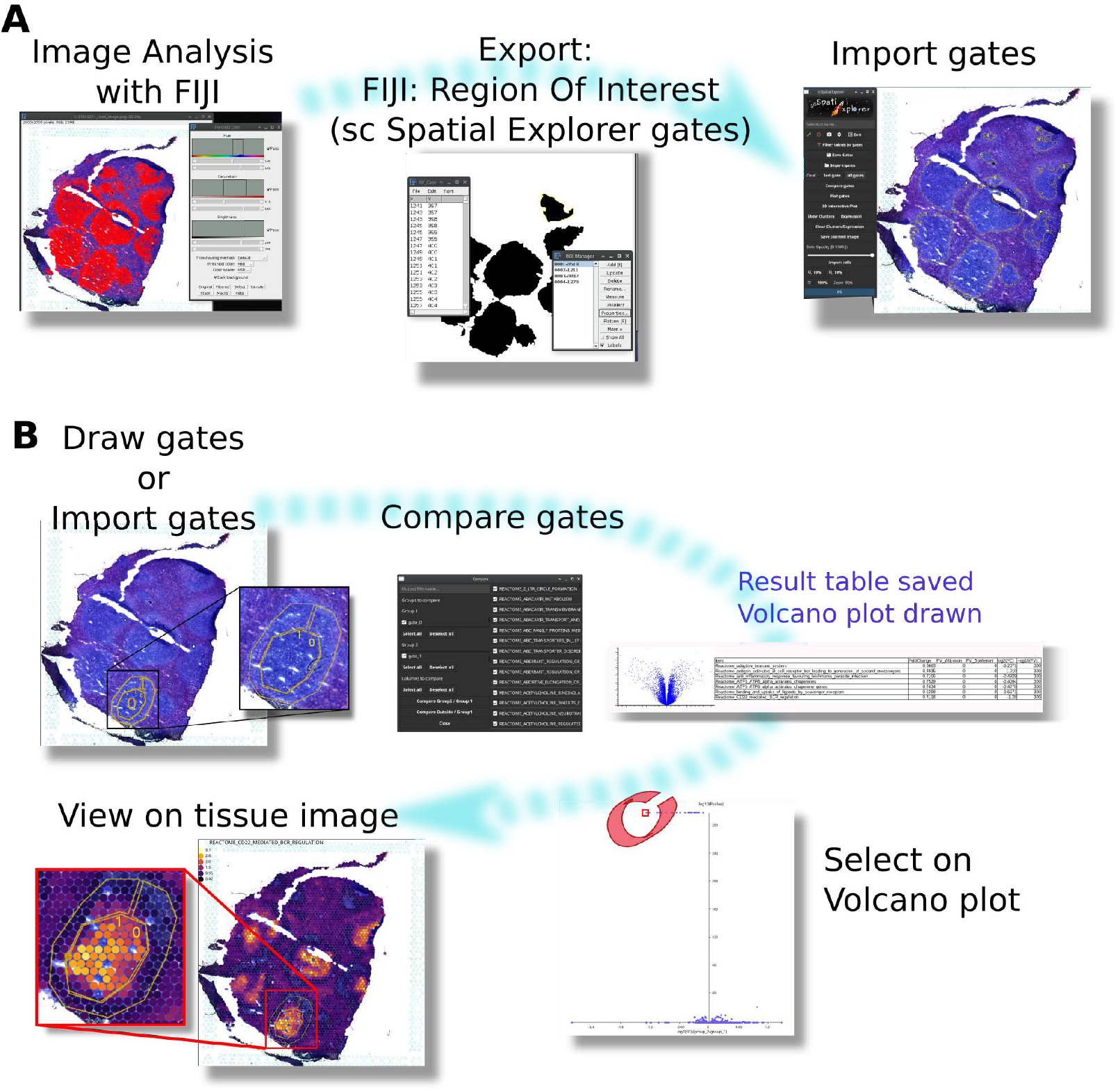
A complex gate pattern was obtained by image analysis. Using ImageJ/Fiji tools, Region Of Interest (ROI, here splenic follicles) were defined, exported and imported in Single-Cell Spatial Explorer (A). Gates previously defined with ImageJ/Fiji were compared in Single-Cell Spatial Explorer on the Reactome pathway (> 2500 pathways) scores computed in Single-Cell Signature Explorer^1^, resulting in a P-value vs fold change table and volcano plot (B). On the volcano plot, a dot was selected and the corresponding pathway (‘CD22-mediated BCR regulation’) visualized on the tissue image.

**Supplementary Figure 2.**
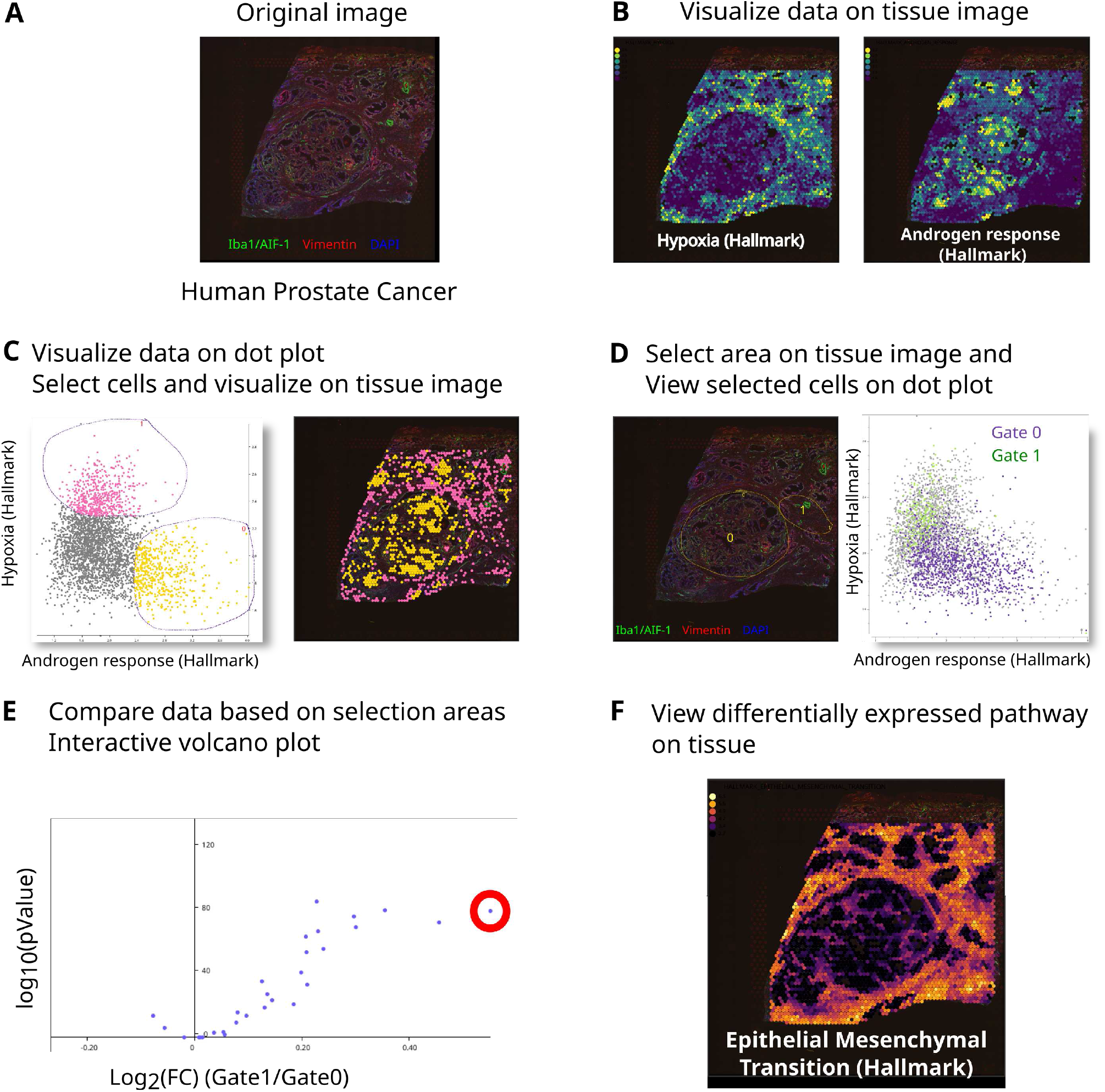
The human prostate cancer dataset was downloaded from 10x Genomics website. [10x Genomics website](https://www.10xgenomics.com/resources/datasets/human-prostate-cancer-adjacent-normal-section-with-if-staining-ffpe-1-standard-1-3-0). The tissue was stained by immunofluorescence using antibodies Iba1/AIF-1 (green),Vimentin (red), DAPI (blue) (A). Data were normalized using Seurat^2^, exported and Hallmark pathways were scored using Single-cell Explorer Scorer^1^. Using Single-Cell Spatial Explorer, scores from hallmark pathways (hypoxia, left; Androgen Response right) are visualized (B). (C) On the “Hypoxia” / “Androgen Response” dot plot, cells were selected (Hypoxia^+^AndrogenResponse^−^ in pink and Hypoxia^−^AndrogenResponse^+^ in yellow) and visualized on the tissue image. (D) Area from the tissue image were selected and cells from this area were visualized on the Hypoxia” / “Androgen Response” dot plot (top) or compared using all the hallmark pathways resulting in a table and a volcano plot (E). Interesting dot can be selected (red circle in E) and data visualized on the tissue image (F).

**Supplementary Figure 3.**
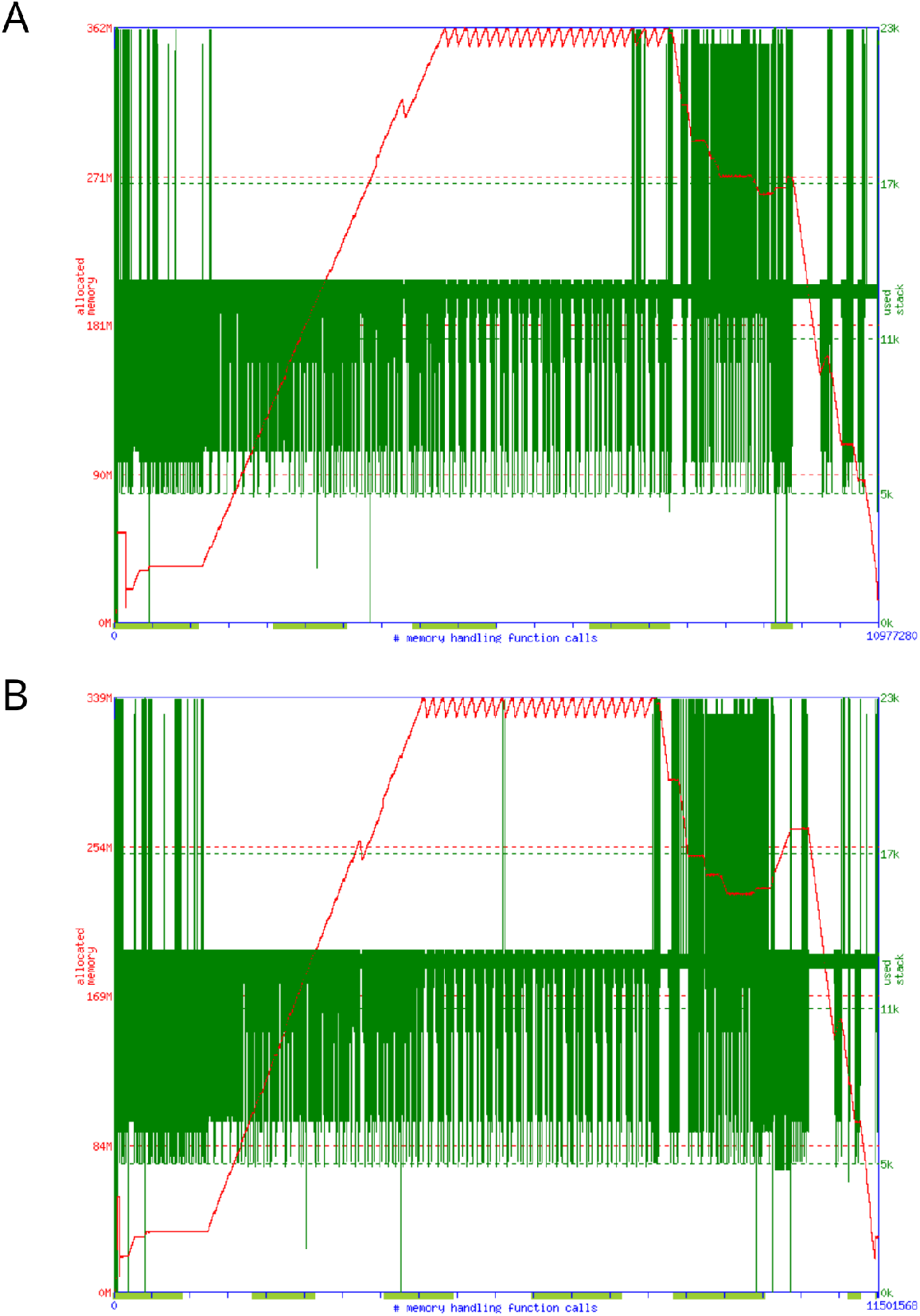
Single-Cell Spatial Explorer memory usage. Output of [memusage](https://www.man7.org/linux/man-pages/man1/memusage.1.html) memory profiling software on a entry level computer. Operating system: GNU Linux Manjaro Xfce, Intel(R) Core(TM) i5-3470 CPU @ 3.20GHz with 8 GB RAM. The session monitored contains a cluster plot, followed by 50 expression plots, a volcano plot and 3 min pause before closing the software. A) Single-Cell Spatial Explorer session with a 2798×202 dataframe, 2000×2000. 362 MB max allocated memory. B) Single-Cell Spatial Explorer session with a 2798×9885 dataframe, 2000×2000 pixels image. 339 MB max allocated memory. Red: allocated memory. Green: stack usage.

**Supplementary Figure 4.**
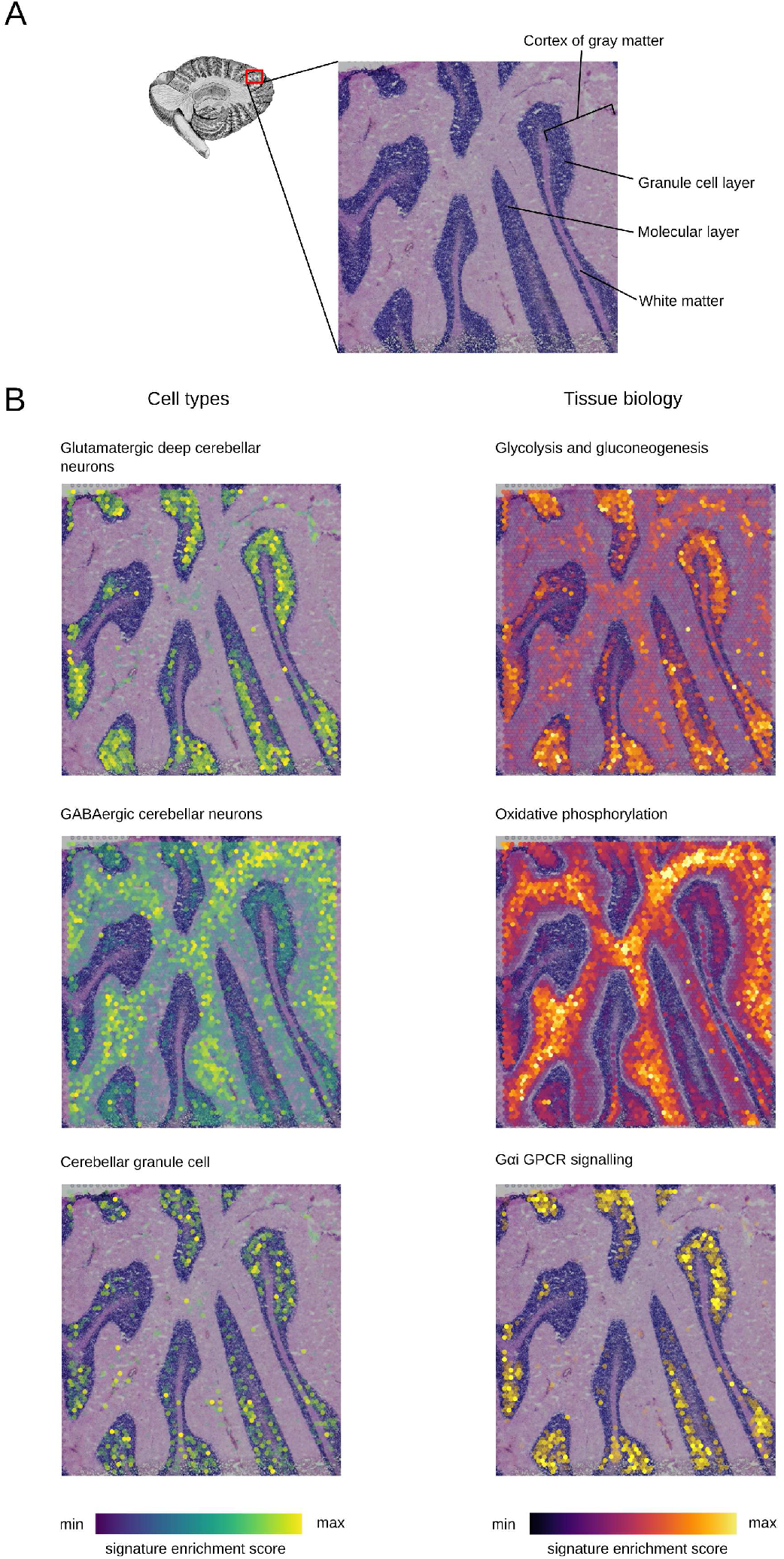
The human cerebellus dataset was downloaded from [10x Genomics website](https://www.10xgenomics.com/resources/datasets/human-cerebellum-whole-transcriptome-analysis-1-standard-1-2-0). The tissue was stained with hematoxylin and aqueous eosin (HE) (A). Data were normalized using Seurat^2^, and pathways from MSIgDB^3^ were scored using Single-cell Explorer Scorer^1^. Using Single-Cell Spatial Explorer, signatures of cell types (Viridis gradient heatmap) and tissue biology (Inferno gradient heatmap) are visualized through a min-max scale and opacity threshold (B).

**Supplementary Figure 5.**
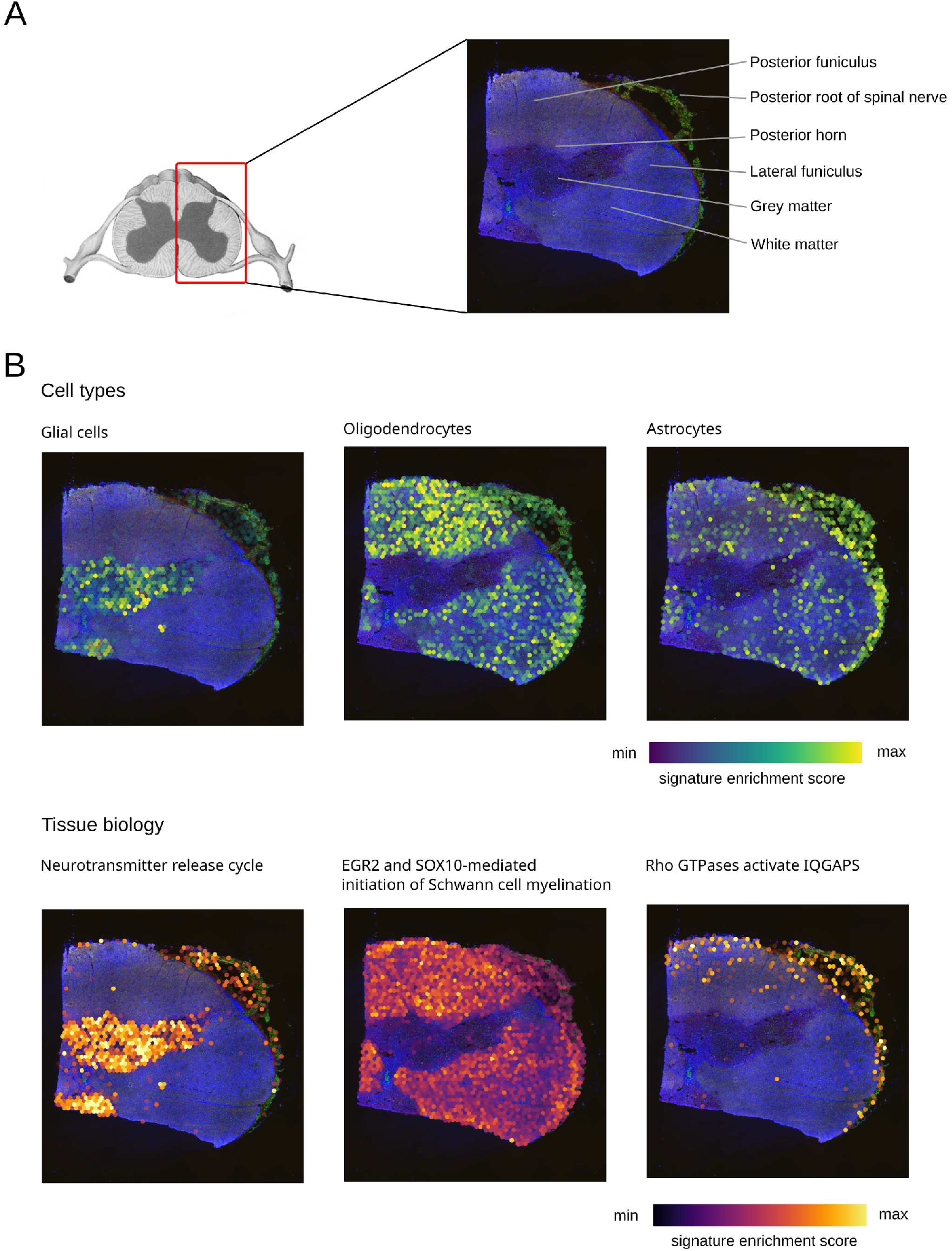
The human spinal cord dataset was downloaded from [10x Genomics website](https://www.10xgenomics.com/resources/datasets/human-spinal-cord-whole-transcriptome-analysis-stains-dapi-anti-snap-25-anti-gfap-anti-myelin-cn-pase-1-standard-1-2-0). The tissue was stained by immunofluorescence using antibodies Anti-SNAP25 (green), Anti-GFAP (pink), Anti-Myelin CNPase (red), DAPI (blue) (A). Data were normalized using Seurat^2^, and pathways from MSIgDB^3^ were scored using Single-cell Explorer Scorer^1^. Using Single-Cell Spatial Explorer, signatures of cell types (Viridis gradient heatmap) and tissue biology (Inferno gradient heatmap) are visualized through a min-max scale and opacity threshold (B).

**Supplementary Figure 6.**
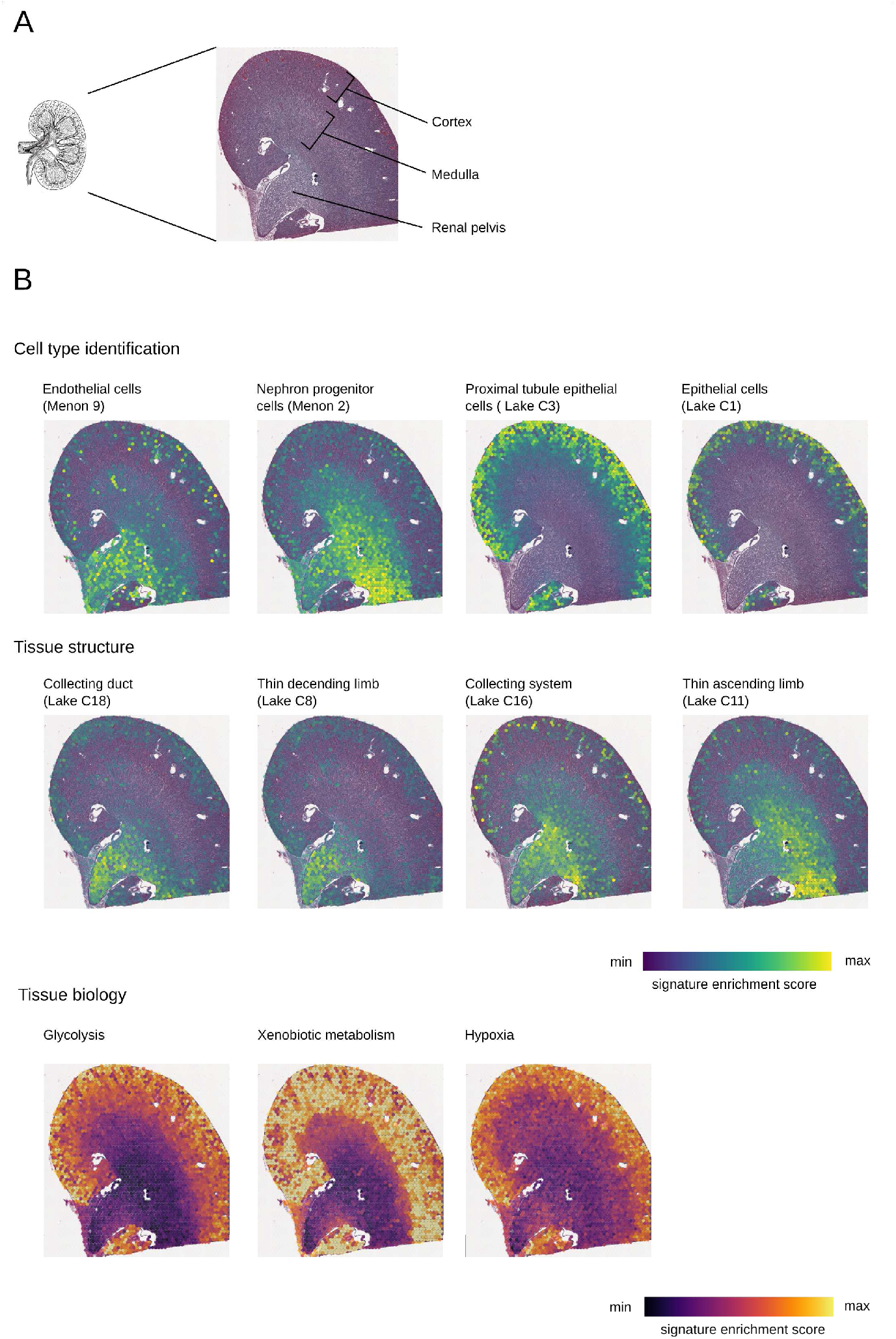
The mouse kidney dataset was downloaded from [10x Genomics website](https://www.10xgenomics.com/resources/datasets/adult-mouse-kidney-ffpe-1-standard-1-3-0). The tissue was stained with hematoxylin and aqueous eosin (HE) (A). Data were normalized using Seurat^2^, and pathways from MSIgDB^3^ were scored using Single-cell Explorer Scorer^1^. Using Single-Cell Spatial Explorer, signatures of cell types, tissue structure (Viridis gradient heatmap) and tissue biology (Inferno gradient heatmap) are visualized through a min-max scale and opacity threshold (B).

**Supplementary Figure 7.**
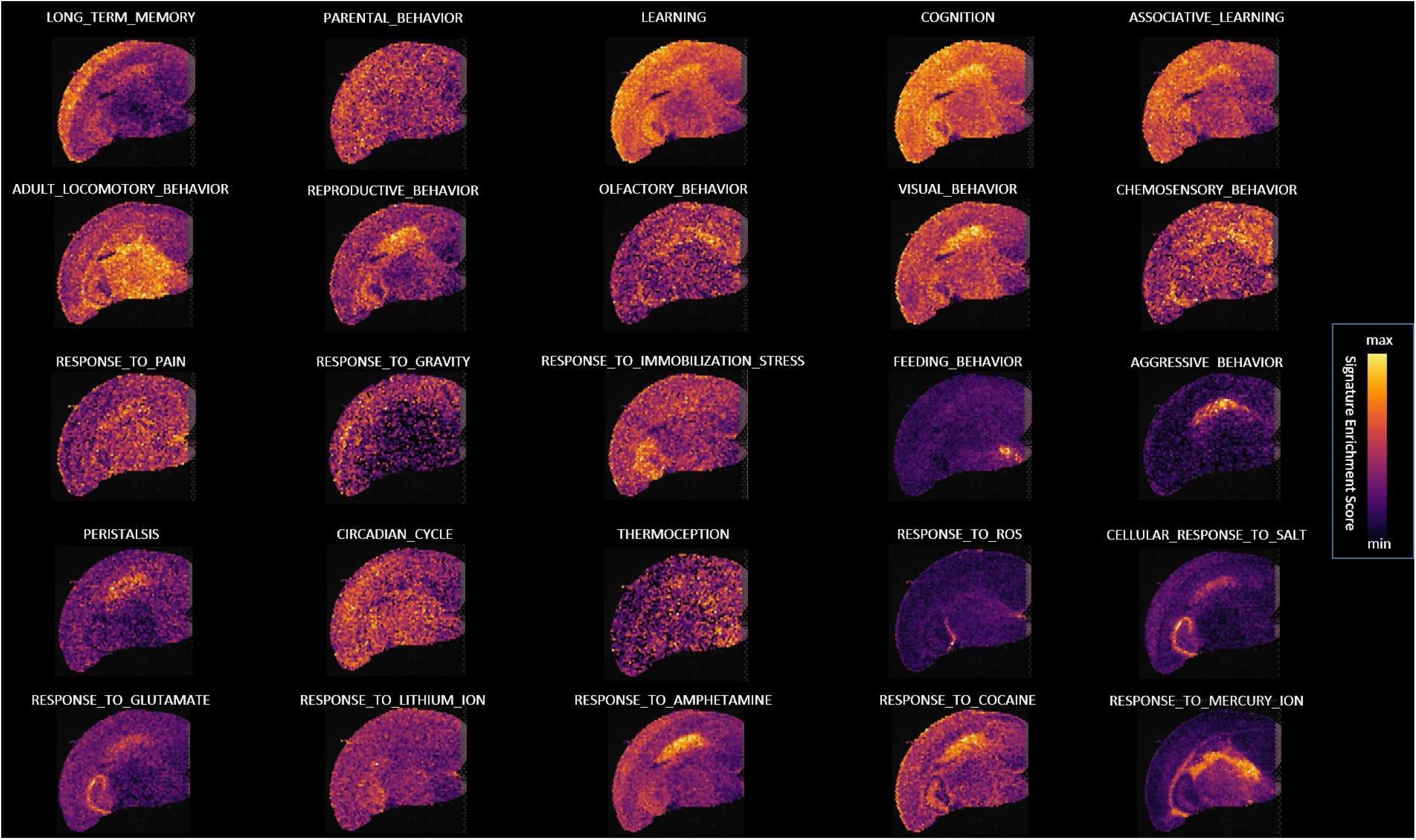
The mouse brain dataset was downloaded from [10x Genomics website](https://www.10xgenomics.com/resources/datasets/adult-mouse-brain-ffpe-1-standard-1-3-0). Data were normalized using Seurat^2^, and signatures from MSIgDB^3^ were scored using Single-cell Explorer Scorer^1^. Using Single-Cell Spatial Explorer, the specified behavioral signatures from GO-BP are visualized (Inferno gradient heatmap) through a min-max scale without opacity threshold.

**Appendix 1 Table1:**
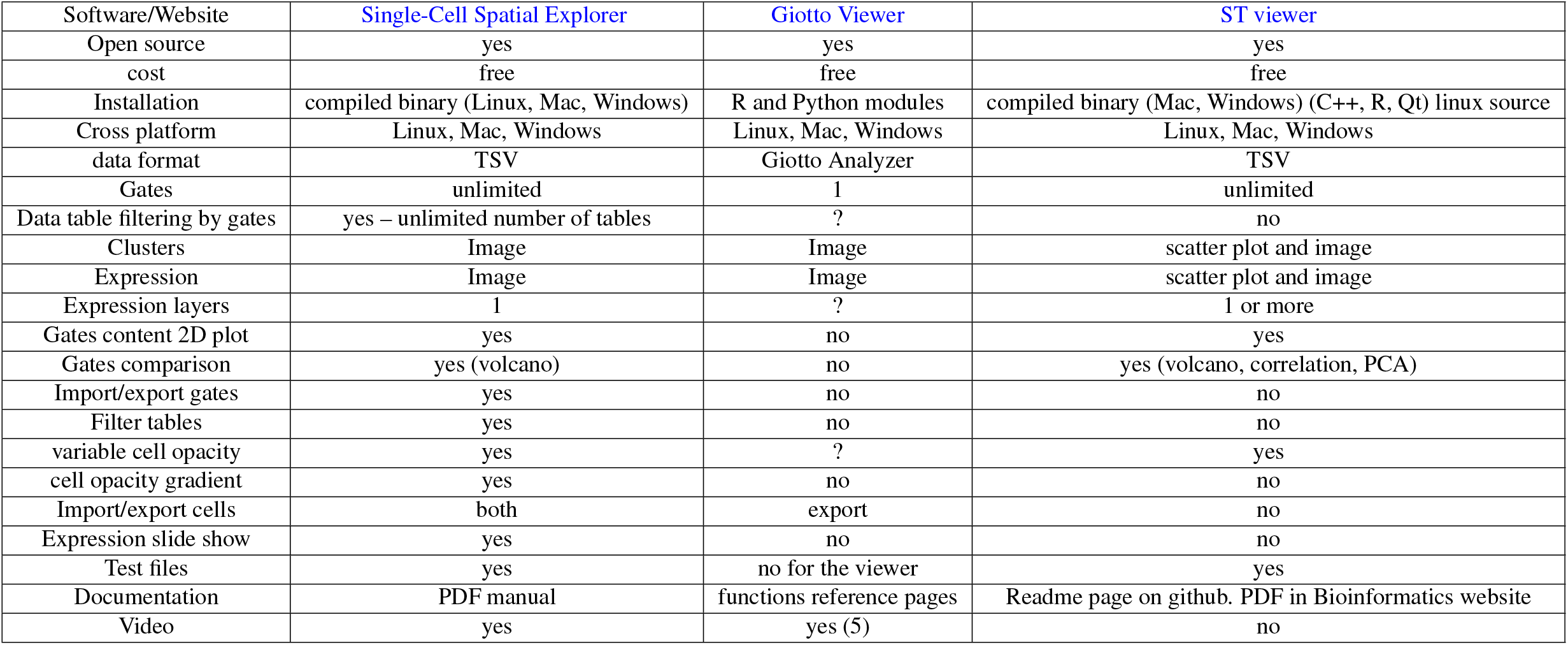
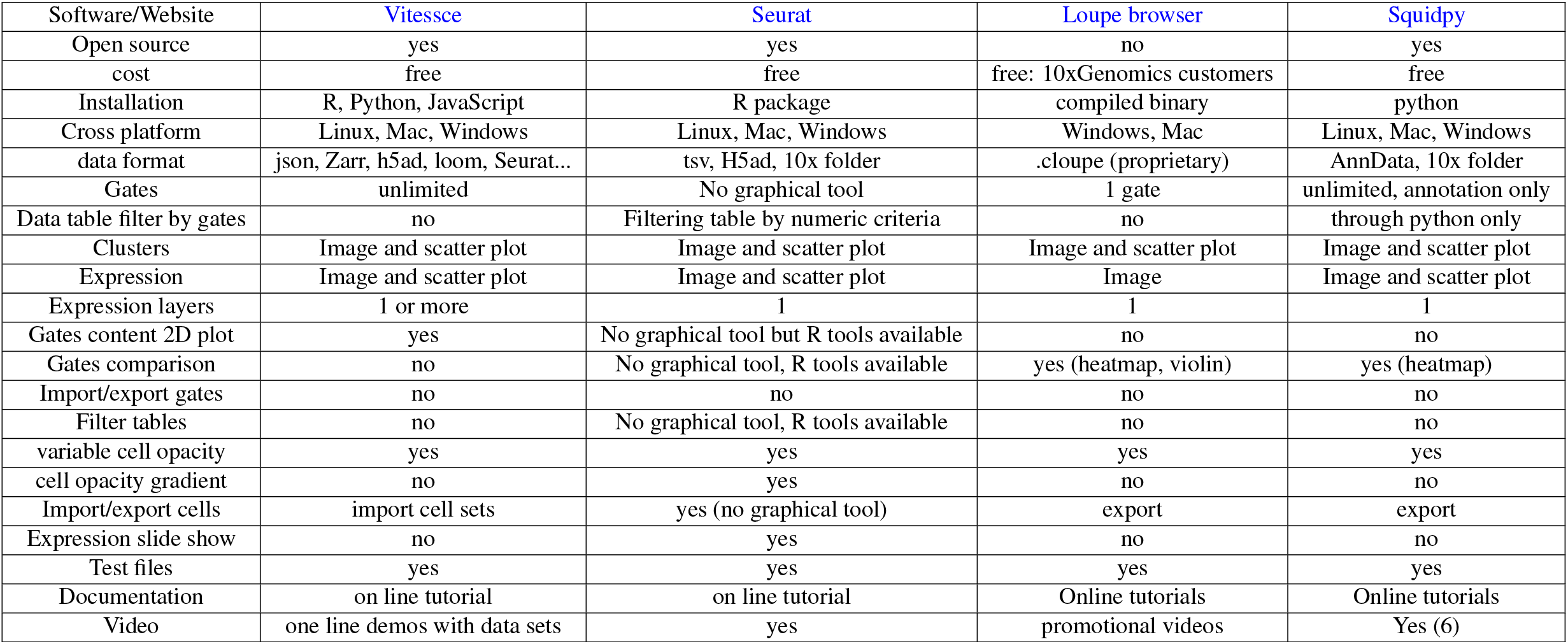
spatial transcriptomics softwares comparison

| Software/Website | Single-Cell Spatial Explorer | Giotto Viewer | ST viewer |
| --- | --- | --- | --- |
| Open source | yes | yes | yes |
| cost | free | free | free |
| Installation | compiled binary (Linux, Mac, Windows) | R and Python modules | compiled binary (Mac, Windows) (C++, R, Qt) linux source |
| Cross platform | Linux, Mac, Windows | Linux, Mac, Windows | Linux, Mac, Windows |
| data format | TSV | Giotto Analyzer | TSV |
| Gates | unlimited | 1 | unlimited |
| Data table filtering by gates | yes – unlimited number of tables | ? | no |
| Clusters | Image | Image | scatter plot and image |
| Expression | Image | Image | scatter plot and image |
| Expression layers | 1 | ? | 1 or more |
| Gates content 2D plot | yes | no | yes |
| Gates comparison | yes (volcano) | no | yes (volcano, correlation, PCA) |
| Import/export gates | yes | no | no |
| Filter tables | yes | no | no |
| variable cell opacity | yes | ? | yes |
| cell opacity gradient | yes | no | no |
| Import/export cells | both | export | no |
| Expression slide show | yes | no | no |
| Test files | yes | no for the viewer | yes |
| Documentation | PDF manual | functions reference pages | Readme page on github. PDF in Bioinformatics website |
| Video | yes | yes (5) | no |

| Software/Website | Vitessce | Seurat | Loupe browser | Squidpy |
| --- | --- | --- | --- | --- |
| Open source | yes | yes | no | yes |
| cost | free | free | free: 10xGenomics customers | free |
| Installation | R, Python, JavaScript | R package | compiled binary | python |
| Cross platform | Linux, Mac, Windows | Linux, Mac, Windows | Windows, Mac | Linux, Mac, Windows |
| data format | json, Zarr, h5ad, loom, Seurat... | tsv, H5ad, 10x folder | .cloupe (proprietary) | AnnData, 10x folder |
| Gates | unlimited | No graphical tool | 1 gate | unlimited, annotation only |
| Data table filter by gates | no | Filtering table by numeric criteria | no | through python only |
| Clusters | Image and scatter plot | Image and scatter plot | Image and scatter plot | Image and scatter plot |
| Expression | Image and scatter plot | Image and scatter plot | Image | Image and scatter plot |
| Expression layers | 1 or more | 1 | 1 | 1 |
| Gates content 2D plot | yes | No graphical tool but R tools available | no | no |
| Gates comparison | no | No graphical tool, R tools available | yes (heatmap, violin) | yes (heatmap) |
| Import/export gates | no | no | no | no |
| Filter tables | no | No graphical tool, R tools available | no | no |
| variable cell opacity | yes | yes | yes | yes |
| cell opacity gradient | no | yes | no | no |
| Import/export cells | import cell sets | yes (no graphical tool) | export | export |
| Expression slide show | no | yes | no | no |
| Test files | yes | yes | yes | yes |
| Documentation | on line tutorial | on line tutorial | Online tutorials | Online tutorials |
| Video | one line demos with data sets | yes | promotional videos | Yes (6) |

### Supplementary discussion

#### 1 Single-Cell Spatial Explorer features

1. Single-Cell Spatial Explorer is distributed with a detailed [PDF manual](https://github.com/FredPont/spatial/blob/main/doc/Manual_Single_Cell_Spatial_Explorer.pdf) and [video tutorials](https://github.com/FredPont/spatial#demo-videos)
2. Single-Cell Spatial Explorer is ready to use in a pre-compiled binary, [no installation required](https://youtu.be/LBBkN_rJHuc).
3. cross-platform (the graphical interface and the software are coded in pure Go)
4. low memory usage
5. compatible with any PNG image associated with any text file with tab separator containing XY coordinates of the image.
6. compatible with any numeric data: gene expression, pathway scores, antibody expression etc…
7. unlimited number of gates.
8. import/export gates in ImageJ/Fiji format.
9. [extract cells and sub-tables](https://youtu.be/rSefd0pmc7g) delimited by the gates on an unlimited number of tables. Exportation is done in TAB separated files for great interoperability.
10. 2D plots of the cells inside the gates with any XY coordinates: t-SNE, UMAP, gene expression, pathway scores, antibody expression etc…
11. interactive 2D plot to show the selected cells on a t-SNE, UMAP or any other coordinates on the image and to filter the data tables into sub-tables.
12. cluster display with 3 color gradients, custom color palette, custom dot opacity and custom dot size. Shuffle color option change color positions on the map leading to almost 2 billions of possible images with 12 clusters.
13. display any kind of cell expression (genes, pathways, antibodies…) with 7 preset gradients, custom legend color, dot opacity and custom dot size. The gradients are simple two colors maps and rainbow colors maps Turbo, Viridis and Inferno to [optimize accuracy and details visualisation](https://ai.googleblog.com/2019/08/turbo-improved-rainbow-colormap-for.html).
14. Min/Max intensity sliders to tune image contrast or remove artefacts due to outliers.
15. Expression opacity gradient with min/max threshold.
16. slide show to review many cell expression maps without need of repetitive click.
17. screenshot or native resolution image exportation.
18. import and display an unlimited number of cells list by repetitive click on the “import cells” button. The format is directly compatible with Single-Cell Virtual Cytometer^4^.
19. compare two groups of gates together in the whole dataset.
20. compare one group of gates against all the remaining cells.
21. draw an interactive volcano plot after gate comparison.
22. plot cell expression of a selected dot in the volcano plot.
23. export volcano plot and the corresponding data table.
24. image zoom 10-200%

#### 2 Memory footprint

Memory usage was a priority in the development of Single-Cell Spatial Explorer. Indeed, spatial transcriptomics is constantly evolving and the resolution and the number of cells increases rapidly. For most functions, Single-Cell Spatial Explorer does not load the whole dataset in RAM but loads only a needed subset. A large data set will increase hard disk usage and slow down the software; but it will still be possible to display it with an adequate amount of RAM. Single-Cell Spatial Explorer RAM footprint analysis with [memusage](https://www.man7.org/linux/man-pages/man1/memusage.1.html) (see supplementary fig. 3) for 2798 × 202 dataframe and 2798 × 9885 dataframe, shows a memory usage < 400*MB*. This memory footprint is very low, so Single-Cell Spatial Explorer does not require a powerful computer at least for the current resolution of spatial transcriptomics.

#### 3 Compatibility with ImageJ/Fiji

This macro was used to export the regions of interest: Macro ImageJ/Fiji to export Region Of Interest (ROI) coordinate from ROI manager:

~~~
for (i=0; i<roiManager(“count”); i++) {
 roiManager(“select", i);
 roiManager(“Set Color", “yellow”);
roiManager(“Set Line Width", 0);
 saveAs(“Results", “MyFolder/XY_
 Coordinates_"+i+".csv”);
}
~~~

#### 4 Comparison with existing softwares

[Seurat](https://satijalab.org/seurat/) ^2^ is a R package designed for single-cell RNA-seq data exploration. It is a R package able to do pre-processing tasks (such as quality controls, data normalisation, samples aggregation and batch effect correction), data analysis (reduce the dimensionality with PCA/t-SNE/UMAP, cluster analysis, differential gene expression, …) and data visualisation. Seurat is a very powerful tool targeting users with programming knowledge such as bio-informaticians with a solid R expertise.

The [Giotto](https://rubd.github.io/Giotto_site/)^5^ package consists of two modules, Giotto Analyzer and Viewer, which provide tools to process, analyze and visualize single-cell spatial expression data. It is compatible with 9 different state-of-the-art spatial technologies, including in situ hybridization (seqFISH+, merFISH, osmFISH), sequencing (Slide-seq, Visium, STARmap) and imaging-based multiplexing/proteomics (CyCIF, MIBI, CODEX). Data analysis is performed first in Giotto Analyzer using command lines and exported then to Giotto viewer. A solid experience in R is needed to install Giotto since many R and Python packages are needed and some dependencies are not listed in the installation note ^1^.

In the Giotto Analyzer, the image alignment should be done by trial and error: the place and the zoom of the image are controlled by numerical parameters. Giotto Analyzer can filtrate genes ans cells for quality controls. At the time of the manuscript, the [link](http://spatial.rc.fas.harvard.edu/giotto-viewer/) to the Giotto viewer was broken and the documentation [“How to switch between Giotto Analyzer and Viewer?”](https://rubd.github.io/Giotto_site/articles/getting_started.html#description) was not available with the status “work in progress”.

[ST viewer](https://github.com/SpatialTranscriptomicsResearch/st_viewer)^6^ is a compiled software to perform analysis and visualization of spatial transcriptomic datasets. The ST viewer enables users to visualize the location of one or multiple genes in real time in a stand-alone desktop application. ST viewer can filter and normalize data with DESeq2, reduce the dimensionality with t-SNE or PCA and compute clusters. A binary for linux is not available. Unfortunately we were not able to test ST viewer on windows since it does not start. This issue might be related to the R version though compatible versions are not documented and remain without answer in dedicated [forum](https://github.com/jfnavarro/st_viewer/issues/38). We opened an [issue](https://github.com/jfnavarro/st_viewer/issues/50) during manuscript preparation. The Mac version need QT recompilation which is not accessible to the average user.

[Vitessce](http://vitessce.io/)^7^ (Visual integration tool for exploration of spatial single cell experiments) is an open-source interactive visualization framework for exploration of multi-modal and spatially-resolved single-cell data; It presents a modular architecture compatible with transcriptomic, proteomic, genome-mapped, and imaging data types. Vitessce can be used as a standalone web application or in Python or R environments. This software has nice online demos with 8 data sets.

Loupe Cell browser is a dedicated visualization and analysis tool for scRNAseq developed for analysing scRNAseq datasets produced by 10xGenomic platforms. It allows importing datasets and visualizing custom projections of either gene expression or antibody-only datasets, across t-SNE or UMAP computed by the Cell Ranger pipeline. Loupe Cell browser also provides Moran’s I for pattern analysis of single gene expression level views across the image. Despite its ease of use however, this tool does not provide signature visualizations, nor the corresponding image pattern analyses.

[SquidPy](https://squidpy.readthedocs.io/en/stable/index.html)^8^ is a python library dedicated to the analysis and visualisation of single-cell spatial transcriptomics datasets. The installation as only been tested on Linux, and requires some basic python knowledge. This library is used after Scanpy QC & normalisation of the spatial dataset. This powerfull tool gives a lot of analysis possibilities, such as cluster annotation, cluster features computation, neighborhood enrichment or even ligand-receptor interaction analysis. This tool also offers the possibility to integrate other libraries to perform complementary analysis. SquidPy has an easy-to-use graphical interface, but main analysis features as described earlier are only available through command line, thus could prevent some users to explore their data to the fullest.

Commercial softwares, such as [BBrowser](https://bioturing.com/bbrowser) or [Partek Flow](https://www.partek.com/partek-flow/) are powerful analysis and visualisation softwares for single cell transcriptome analysis and spatial transcriptomics. However, since they are proprietary, not open source, and their cost is not specified in their website, they could not be tested here.

So currently, quite a few existing open source tools allow to visualize single cell spatial transcriptomics data (see software comparison in supplementary Table 1). We found two softwares, Loupe and STviewer, which are free of charge^2^ and provide a ready to use binary executable like Single-Cell Spatial Explorer. Unfortunately we had issues with the compiled binaries of STviewer and Loupe interoperability is very far from what is possible with Single-Cell Spatial Explorer. By being mainly dedicated to data visualization, Single-Cell Spatial Explorer is mostly recommended for biologists, pathologists and biomedical users that are not familiar with R or command line tools. Single-Cell Spatial Explorer is currently the best alternative for users who want a software which works out of the box, without tedious installation.

## Footnotes

1 such as openssl, libcurl4-openssl-dev, libxml2-dev, libssl-dev, libgmp3-dev, libgsl-dev, libmagick++-dev

2 Loupe is free of charge for 10x Genomics customers.

## References

1. A. Butler, P. Hoffman, P. Smibert, E. Papalexi, and R. Satija. Integrating single-cell transcriptomic data across different conditions, technologies, and species. Nature biotechnology, 36:411–420, June 2018.

2. J.-P. Cerapio, M. Perrier, C.-C. Balança, P. Gravelle, F. Pont, C. Devaud, D.-M. Franchini, V. Féliu, M. Tosolini, C. Valle, et al. Phased differentiation of γ δ t and t cd8 tumor-infiltrating lymphocytes revealed by single-cell transcriptomics of human cancers. Oncoimmunology, 10(1):1939518, 2021.

3. T. J. Chen and N. Kotecha. Cytobank: providing an analytics platform for community cytometry data analysis and collaboration. pages 127–157. Springer, 2014.

4. C.-S. Cho, J. Xi, Y. Si, S.-R. Park, J.-E. Hsu, M. Kim, G. Jun, H. M. Kang, and J. H. Lee. Microscopic examination of spatial transcriptome using seq-scope. Cell, 184(13):3559–3572, 2021.

5. A. Liberzon, C. Birger, H. Thorvaldsdóttir, M. Ghandi, J. P. Mesirov, and P. Tamayo. The molecular signatures database (msigdb) hallmark gene set collection. Cell systems, 1:417–425, Dec. 2015.

6. F. Pont, M. Tosolini, and J.-J. Fournié. Single-Cell Signature Explorer for comprehensive visualization of single cell signatures across scRNA-seq datasets. Nucleic acids research, 47(21):e133–e133, 07 2019.

7. F. Pont, M. Tosolini, Q. Gao, M. Perrier, M. Madrid-Mencía, T. S. Huang, P. Neuvial, M. Ayyoub, K. Nazor, and J.-J. Fournié. Single-cell virtual cytometer allows user-friendly and versatile analysis and visualization of multimodal single cell rnaseq datasets. NAR genomics and bioinformatics, 2(2):qaa025, 2020.

8. S. G. Rodriques, R. R. Stickels, A. Goeva, C. A. Martin, E. Murray, C. R. Vanderburg, J. Welch, L. M. Chen, F. Chen, and E. Z. Macosko. Slide-seq: A scalable technology for measuring genome-wide expression at high spatial resolution. Science, 363(6434):1463–1467, 2019.

9. J. Schindelin, I. Arganda-Carreras, E. Frise, V. Kaynig, M. Longair, T. Pietzsch, S. Preibisch, C. Rueden, S. Saalfeld, B. Schmid, et al. Fiji: an open-source platform for biological-image analysis. Nature methods, 9(7):676–682, 2012.

10. C. A. Schneider, W. S. Rasband, and K. W. Eliceiri. Nih image to imagej: 25 years of image analysis. Nature methods, 9(7):671–675, 2012.

11. P. L. Ståhl, F. Salmén, S. Vickovic, A. Lundmark, J. F. Navarro, J. Magnusson, S. Giacomello, M. Asp, J. O. Westholm, M. Huss, et al. Visualization and analysis of gene expression in tissue sections by spatial transcriptomics. Science, 353(6294):78–82, 2016.

12. R. R. Stickels, E. Murray, P. Kumar, J. Li, J. L. Marshall, D. J. Di Bella, P. Arlotta, E. Z. Macosko, and F. Chen. Highly sensitive spatial transcriptomics at near-cellular resolution with slide-seqv2. Nature biotechnology, 39(3):313–319, 2021.

13. M. Stoeckius, C. Hafemeister, W. Stephenson, B. Houck-Loomis, P. K. Chattopadhyay, H. Swerdlow, R. Satija, and P. Smibert. Simultaneous epitope and transcriptome measurement in single cells. Nature methods, 14(9):865, 2017.

14. T. Stuart, A. Butler, P. Hoffman, C. Hafemeister, E. Papalexi, W. M. Mauck III, Y. Hao, M. Stoeckius, P. Smibert, and R. Satija. Comprehensive integration of single-cell data. Cell, 2019.

15. S. Vickovic, G. Eraslan, F. Salmén, J. Klughammer, L. Stenbeck, D. Schapiro, T. Äijö, R. Bonneau, L. Bergenstråhle, J. F. Navarro, et al. High-definition spatial transcriptomics for in situ tissue profiling. Nature methods, 16(10):987–990, 2019.

16. L. Walker, Z. Cang, H. Ren, E. Bourgain-Chang, and Q. Nie. Deciphering tissue structure and function using spatial transcriptomics. Communications Biology, 5(1):1–10, 2022.

17. F. A. Wolf, P. Angerer, and F. J. Theis. Scanpy: large-scale single-cell gene expression data analysis. Genome biology, 19(1):1–5, 2018.

## References

1. Pont, F., Tosolini, M. & Fournié, J.-J. Single-Cell Signature Explorer for comprehensive visualization of single cell signatures across scRNA-seq datasets. Nucleic acids research 47, e133–e133, DOI: 10.1093/nar/gkz601 (2019).

2. Butler, A., Hoffman, P., Smibert, P., Papalexi, E. & Satija, R. Integrating single-cell transcriptomic data across different conditions, technologies, and species. Nat. biotechnology 36, 411–420, DOI: 10.1038/nbt.4096 (2018).

3. Liberzon, A. et al. The molecular signatures database (msigdb) hallmark gene set collection. Cell systems 1, 417–425, DOI: 10.1016/j.cels.2015.12.004 (2015).

4. Pont, F. et al. Single-cell virtual cytometer allows user-friendly and versatile analysis and visualization of multimodal single cell rnaseq datasets. NAR genomics bioinformatics 2, lqaa025 (2020).

5. Dries, R. et al. Giotto, a toolbox for integrative analysis and visualization of spatial expression data. bioRxiv DOI: 10.1101/701680 (2020).

6. Fernández Navarro, J., Lundeberg, J. & Ståhl, P. L. ST viewer: a tool for analysis and visualization of spatial transcriptomics datasets. Bioinformatics 35, 1058–1060, DOI: 10.1093/bioinformatics/bty714 (2018).

7. Keller, M. S. et al. Vitessce: a framework for integrative visualization of multi-modal and spatially-resolved single-cell data. OSF Prepr. DOI: 10.31219/osf.io/y8thv (2021).

8. Palla, G. et al. Squidpy: a scalable framework for spatial omics analysis. Nat. Methods 19, 171–178, DOI: 10.1038/s41592-021-01358-2 (2022).

